# Antibacterial effects of propolis and brood comb extracts on the causative agent of European Foulbrood (*Melissococcus plutonius*) in honey bees (*Apis mellifera*)

**DOI:** 10.1101/2022.02.21.481376

**Authors:** Stephanie K. Murray, Colin M. Kurkul, Andrew J. Mularo, Vanessa L. Hale, Rachelle M. M. Adams, Reed M. Johnson

## Abstract

Among a long list of parasites and pathogens that threaten the European honey bee (*Apis mellifera*), European Foulbrood (EFB) has become an urgent apiary disease, as epidemic outbreaks are becoming increasingly common. EFB is a bacterial disease of larval honey bees, caused by a gram-positive, anaerobic bacterium, *Melissococcus plutonius*. The most effective current treatment for EFB, oxytetracycline hydrochloride, can disrupt the bee microbiome, cause bee mortality and residues may persist in honey harvested for human consumption. In this study, we explore the efficacy of more sustainable bee-derived solutions, including propolis, honey comb and brood comb ethanol extracts. Using a series of dilutions of these extracts, we determined the minimum inhibitory concertation (MIC) of each bee-derived product on *M. plutonius*, as well as two model bacterial species, *Staphylococcus saprophyticus* (gram-positive) and *Escherichia coli* (gram-negative). Overall, we found that propolis extract was most effective at inhibiting growth of gram-positive bacteria, and that *M. plutonious* was also susceptible to honey comb (MIC = 16.00 mg/mL) and brood comb (MIC = 45.33 mg/mL) extracts, but at much higher concentrations than that of propolis (MIC = 1.14 mg/mL).

## Introduction

The social life history of the European honey bee (*Apis mellifera*) is highly conducive to the spread of pathogens, such as foulbrood (i.e., bacterial disease of the larvae). Both American Foulbrood and European Foulbrood diseases, named for the regions in which they were initially discovered (Milbrath, 2021), are present in honey bee colonies worldwide. American Foulbrood (AFB) is typically more devastating, as the causative agent *Paenibacillus larvae* is spore-forming, and spores are known to remain viable on beekeeping equipment for at least 35 years (Haseman, 1961). While antibiotics such as oxytetracycline hydrochloride (OTC) can be effective against *P. larvae* in the vegetative stage, they are ineffective against spores (Lodesani & Costa, 2005). Additionally, tetracycline antibiotics decrease core microbiota of the honey bee gut, resulting in increased susceptibility to opportunistic bacteria and elevated mortality (Raymann et al., 2017; Soares et al., 2021). These changes have the potential to cause long-term effects on colony fitness (Bulson et al., 2021). Because antibiotics may simply mask AFB infection in a hive (Lodesani & Costa, 2005), colonies diagnosed with AFB are often killed and hive equipment destroyed by fire (Hansen & Brødsgaard, 1999), resulting in a substantial economic loss for beekeepers (estimated at 5 million USD/year; Eischen et al., 2005). In contrast, European Foulbrood (EFB) is caused by *Melissococcus plutonius*, a non-spore-forming bacterium (Bailey, 1957), and is treatable with OTC (Forsgren, 2010). Still, treatment with OTC can be economically damaging as antibiotic residues can prevent beekeepers from selling honey (Gilliam & Argauer, 1981; Matsuka & Nakamura, 1990; Milbrath, 2021; Sporns et al., 1986; Wilson, 1974). Additionally, EFB can easily return after OTC treatment, and outbreaks have become increasingly prominent worldwide (Grossar et al., 2020; Simone-Finstrom & Spivak, 2010).

EFB infection causes discoloration and loss of internal body pressure in affected larvae, leaving them yellow-brown and flaccid (Forsgren, 2010). *Melissococcus plutonius* is a gram-positive, microaerophilic to anaerobic bacterium that thrives in the larval honey bee midgut (Bailey, 1983; Bailey & Collins, 1982; Forsgren, 2010). Infected larvae pass on the bacterium by either defecating or dying within their wax cells, and bacteria are then spread to healthy larvae by adult nurse bees through larval care activities (Bailey, 1983), often resulting in the spread of EFB within a colony and throughout an apiary.

Propolis is a substance bees make by mixing resinous plant exudates with saliva and beeswax (Simone-Finstrom & Spivak, 2010). Bees use propolis to seal cracks in the hive body, reduce the hive entrance, and line the rims of wax cells (Evans & Spivak, 2010; Seeley & Morse, 1976; Strehle et al., 2003). In a natural setting, feral colonies create a “propolis envelope” around the entire surface of their rough nesting cavity (e.g., a tree cavity; Seeley & Morse, 1976), but this behavior is exhibited much less inside hive boxes made from finely milled lumber. Instead, propolis is frequently deposited between boxes, hive covers, and man-made frames that hold beeswax combs. Due to this meticulous behavior, propolis can be easily harvested for human use in folk medicine (Marcucci, 1995; Przybyłek & Karpiński, 2019). Propolis contains antimicrobial compounds, such as flavonoids, phenolics, terpenoids and aromatic acids, that are present in the source plant exudates (Marcucci, 1995; Przybyłek & Karpiński, 2019). These compounds vary greatly depending on hive location and nearby plants from which the bees collect material (Huang et al., 2014; Sforcin, 2016). For example, propolis in temperature regions originates mostly from *Populus* species (Anđelković et al., 2017; Sforcin, 2016), and thus propolis samples collected in temperate regions (e.g., Ohio) largely contain flavonoids and phenolics (Johnson et al., 1994).

Propolis has been shown to combat various pathogenic threats to bees, including fungal parasites like *Nosema* spp. and *Ascosphaera apis*, the causative agents of Nosemosis and larval chalkbrood disease, respectively (Borba et al., 2015; Simone-Finstrom & Spivak, 2012). It has also demonstrated effects against bee viral infections, including deformed wing virus and black queen cell virus (Borba et al., 2015; Drescher et al., 2017). Additionally, bees from hives that were supplemented with additional propolis were found to exhibit lower expression of immune genes (Simone et al., 2009).

With a recent increase in EFB outbreaks and the need for more sustainable treatments, we aimed to test the antibacterial properties of extracts from propolis and other bee-derived products (i.e., wax brood combs and honey combs) against *M. plutonius*. We are interested in the effects of wax combs previously used for brood rearing on honey bee health, due to the deposition of both antimicrobial substances (i.e., propolis and honey; Pasupuleti et al., 2017) and bee frass over time. Bees use propolis to increase the structural integrity of all wax combs (Strehle et al., 2003), but brood comb darkens in color and gradually becomes a composite material, as the wax accumulates larval silk, castings, and frass with repeated cycles of brood rearing (Hepburn & Kurstjens, 1988). Due to this build-up of waste materials, as well as the potential for build-up of pesticide residues, beekeepers are often encouraged to replace their used brood comb after several years (Jaycox, 1979) However, the presence and potential benefits of propolis in wax cells is a relatively unexplored research topic. Surveys of beekeepers indicate that older brood comb may be associated with improved winter survival (https://research.beeinformed.org/survey/), possibly due to the propolis constituents or other components accumulated by wax through repeated brood-rearing cycles.

In the present study, we tested the antibacterial properties of ethanol extracts of (1) propolis, (2) brood comb (wax comb in which immature bees had previously been reared), and (3) honey comb (wax comb used exclusively for honey storage) against *M. plutonious*, the bacterial agent causing EFB. Additionally, these extracts were tested against two model bacteria, gram-negative *Escherichia coli* and gram-positive *Staphylococcus saprophyticus*. The latter species are not associated with honey bee disease; however, they are pathogenic bacteria in mammals and are frequently used in antimicrobial testing. Historically, propolis extracts have not demonstrated substantial inhibition of gram-negative bacteria (Przybyłek & Karpiński, 2019), therefore, *E. coli* was included to serve as a negative control. We measured the optical density of all bacterial cultures after exposure to propolis, brood comb and honey comb extracts to determine the lowest concentration, or minimum inhibitory concentration (MIC), that inhibits bacterial growth.

## Materials and Methods

### Colony setup

A total of six experimental colonies were established in standard eight-frame Langstroth-style hive equipment at three apiary locations, with two colonies at each site, in spring of 2018. Apiaries were located at (1) the Waterman Agriculture and Natural Resources Laboratory in Columbus, Ohio (40.0104° N, 83.0400° W), (2) the Ohio Agricultural Research and Development Center in Wooster, Ohio (40.7818° N, 81.9305° W), and (3) the Muck Crops Agricultural Research Station in Willard, Ohio (41.010200° N, 82.731350° W). New waxed black plastic foundation (9FDBW, Pierco) in new wooden frames were fitted in either newly constructed or thoroughly cleaned deep hive boxes (KD-603, Mann Lake). A new or thoroughly cleaned metal queen excluder (HD-121, Mann Lake) was used to restrict the queen and rearing of immature bees, or brood, to the bottom box so that the upper box could only be used for honey storage (Fig. 1).

**Figure 1.**
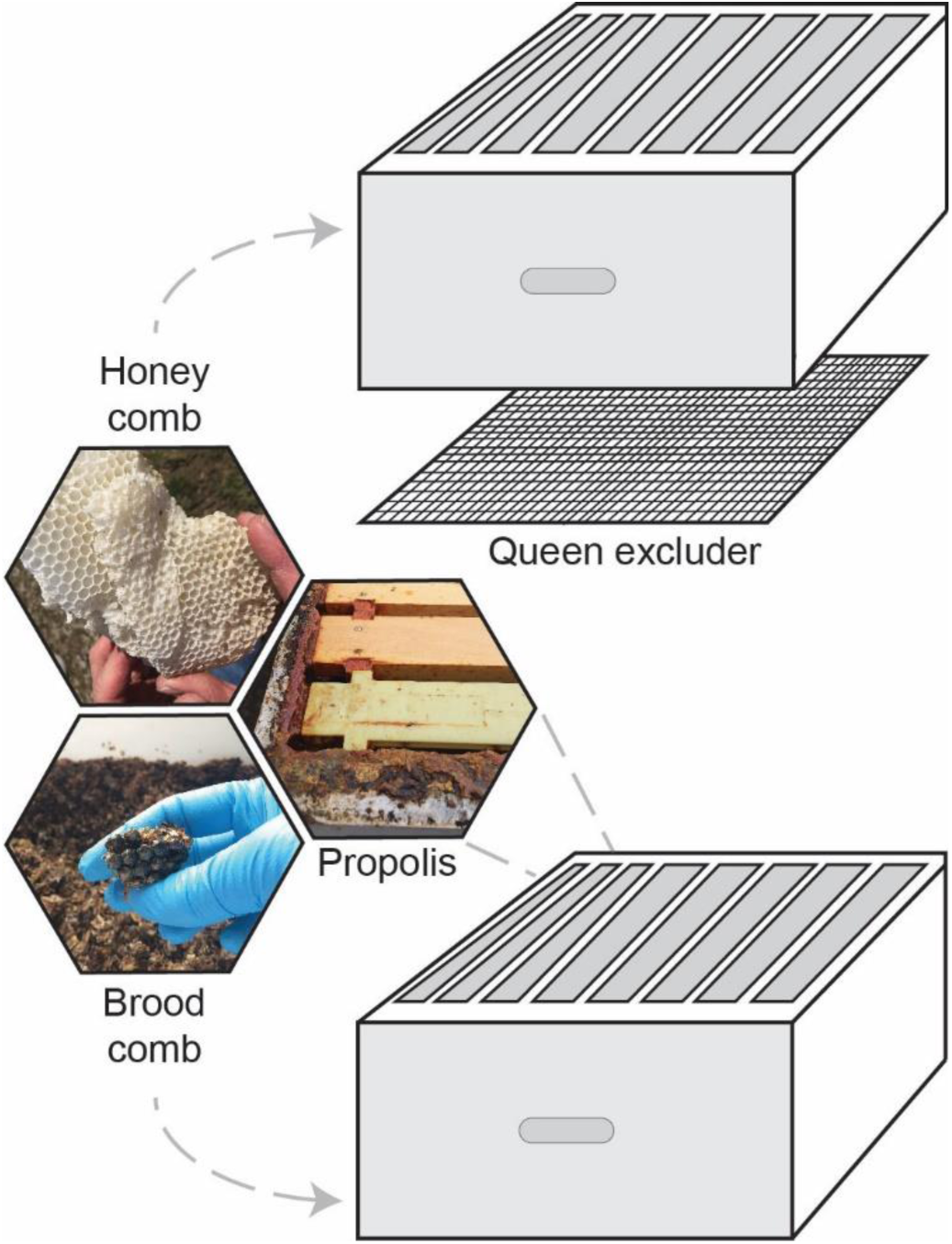
Experimental colony design. Two eight-frame deep boxes separated by a queen excluder, to restrict the queen, brood and brood comb to the lower box and allow construction of comb for honey storage in the upper box. Propolis was collected from hive crevices.

### Wax and propolis collection and extraction

Using a freshly cleaned hive tool, empty combs (free of honey or brood) and propolis samples were collected from each site in late-winter of 2019 (approximately ten months later) and stored in darkness at −20°C until extraction. Before comb samples or propolis were extracted, they were first ground to a fine powder using liquid nitrogen and a food processor (FP2500B, Black & Decker). Separate food processors were used for each sample type to prevent cross contamination. Two grams of each powdered sample were placed in 20 mL of 95% ethanol (1:10 w/v) and sonicated (HB-S-49DHT, Kenda) for 60 minutes at up to 60°C. The resulting extract was gravity filtered using grade one filter paper to remove particulate matter, evaporated under a stream of nitrogen at 20°C, and resuspended in 10 mL of 70% ethanol. Ethanol was chosen as it is a commonly used solvent in propolis research and many of the antimicrobial compounds present are soluble in it. In total, nine crude ethanol extracts were made, three each of honey comb, brood comb and propolis corresponding to the three apiary locations. Concentrations were measured after evaporating crude extracts. The average concentration of dry material in crude extracts was 16.0 mg/mL for honey comb, 45.33 mg/mL for brood comb, and 72.83 mg/mL for propolis. A series of eight two-fold dilutions were then made for testing against bacteria. Diluted extract concentrations are listed in Fig. 2. All extracts were stored in darkness at 4°C.

**Figure 2.**
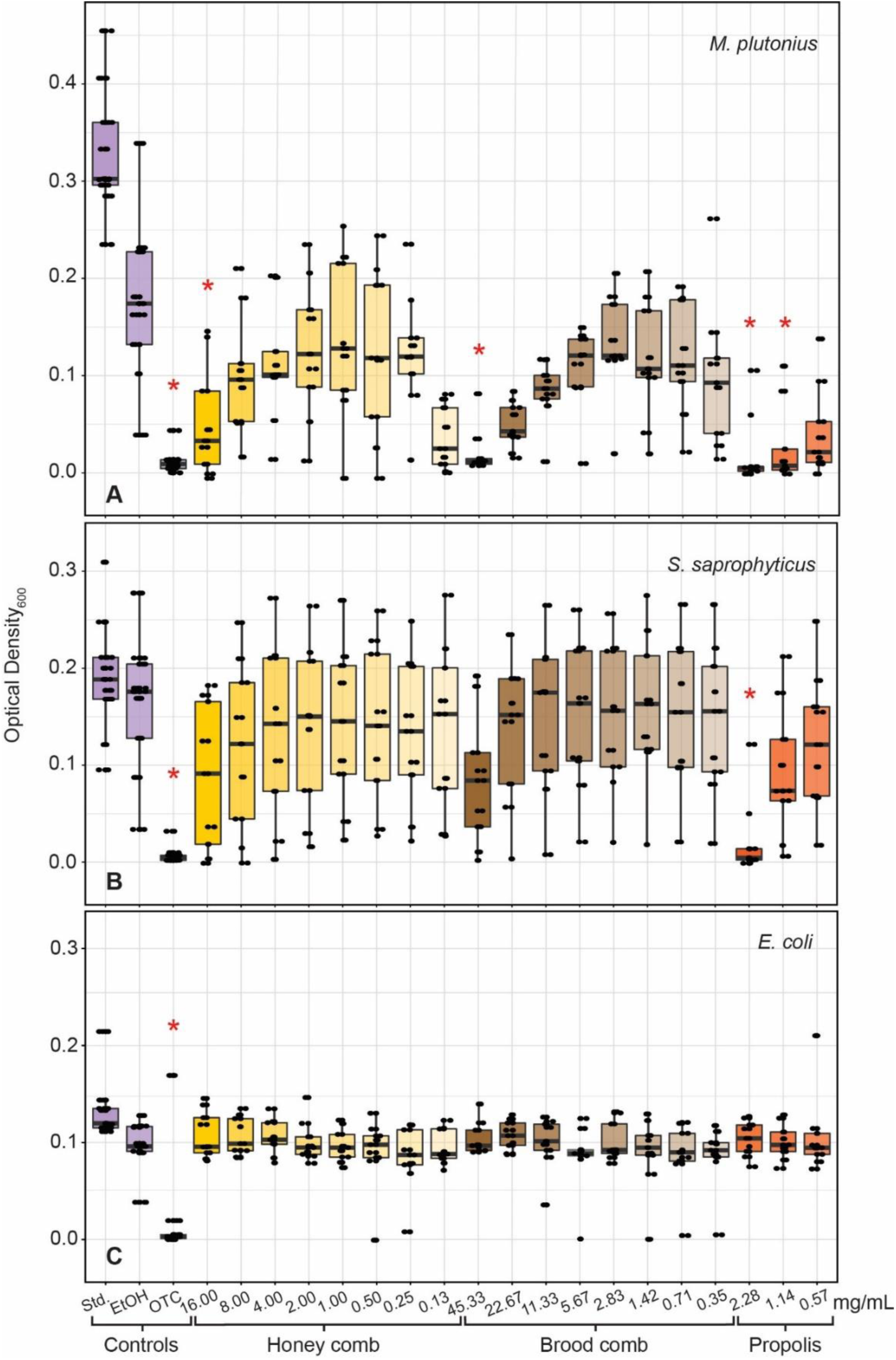
OD_600_ of (A) *M. plutonius*, (B) *S. saprophyticus* and (C) *E. coli* at mid-log phase when treated with a two-fold dilution series of 70% ethanol extracts of honey comb, brood comb and propolis. Comb and propolis were collected from three locations, with three technical replicates each (n=9). Ethanol (70%) was used as a solvent control (EtOH) and Oxytetracycline hydrochloride (1mg/mL) was used as a positive control (OTC). Untreated wells were used as a standard growth control (Std.). MICs and controls significantly different (p<0.05, Dunn’s Multiple Comparison test with Bonferonni correction) from solvent control are indicated with asterisks (*).

### Bacterial Cultures

*Melissococcus plutonius* was isolated from EFB-infected larvae collected at the Ohio State University Wooster Campus in spring 2018. This microaerophilic-anaerobic bacterium was isolated and cultured by inoculating modified basal medium (Forsgren et al., 2013) with fresh infected larvae that were ground with a sterile pestle in 1x PBS solution. Growth medium (Table S1) contained yeast extract, glucose, sucrose, L-cysteine, 1M KH_2_PO_4_ and 2.5M KOH (AC400405000, D16-500 and P285-500, Fisher Scientific; P5958 and W326305, Sigma Aldrich). To prevent the growth of secondary bacteria, nalidixic acid was dissolved in 0.1M NaOH (AAB2509606 and S318-100, Fisher Scientific), filter sterilized, and added to the medium at a concentration of 3 μg/ml. *Melissococcus plutonius* was cultured in a vinyl anaerobic chamber (Coy Laboratory Products, Grass Lake, MI) at 37°C (Table S1). Sequencing using primers from Haynes et al. (2013) confirmed the identity of bacteria in culture (GenBank accession: MN886237). Cultures were stored in 50% glycerol for long-term storage at −80°C. Strains of two aerobic bacterial species, *E. coli* (CSH36) and *S. saprophyticus* (ATCC15305), were obtained from collections of the Department of Microbiology at The Ohio State University and cultured using standard protocols in Mueller-Hinton media (OXCM0405B, Fisher Scientific) at 37°C.

### Bacterial Growth Inhibition Assays

We determined the MIC of each treatment using a broth microdilution method modeled after the Clinical and Laboratory Standards Institute (Cockerill et al., 2012) as in Sozanski et al. (2020). Before testing the inhibitory properties of each extract dilution series, bacterial isolates were first inoculated in broth media appropriate for each bacterium (aerobic Meuller-Hinton media or anaerobic *M. plutonius* media) and grown overnight at 37ºC under shaking aerobic conditions (*E. coli* and *S. saprophyticus*) or still anaerobic conditions (*M. plutonius*). To inoculate microplate test wells (96-well flat-bottomed microplates, Thermo Scientific™ 243656) at 5 × 10^5^ CFU/mL, we first diluted overnight cultures to a 0.5 McFarland standard (1.0 × 10^8^ CFU/mL) and further diluted cultures by a factor of five. We then added these diluted cultures to plate wells containing 5 μL of either antimicrobial treatments or controls and 190 μL of appropriate broth media, resulting in a final concentration of 5 × 10^5^ CFU/mL. Each column of wells on the 96-well plate contained the series of two-fold diluted propolis, brood comb or honey comb extracts, with column one containing the most concentrated treatment and column eight containing the most dilute treatment. Columns nine and 10 contained a positive oxytetracycline (OTC) control (1 mg/mL) and a negative solvent control (70% ethanol), respectively. The final two columns on the plate served as a standard growth control and a contamination control. Standard growth control wells contained 195 μL of appropriate broth with 5 μL of diluted culture. Finally, contamination control wells contained 200 μL of appropriate broth. Each 96-well plate was considered one technical replicate. For every extract and bacterium tested, there were three biological replicates, corresponding to the three different source apiaries, and three technical replicates (n = 9).

To measure bacterial growth, plates were placed in an ELx808i™ Absorbance Microplate Reader (BioTek Instruments) with incubation at 37°C. Plates were shaken and an absorbance reading was made at 600 nm (OD_600_) every five minutes for 12 hours (Quigley, 2008). Standard growth controls for each plate were used to determine the mid-log phase of bacterial growth, allowing MIC determination at that time point for each treatment.

### Statistical Analysis and MIC Determination

We first analyzed standard growth wells to determine the mid-log phase of each plate using the R package Growthcurver v3.6.0 (Sprouffske & Wagner, 2016). We recorded OD_600_ readings for bacterial growth in all wells at their corresponding mid-log phase (supplemental data are openly available in Figshare at https://doi.org/10.6084/m9.figshare.19387442) and tested each plate for growth differences among treatments using a Kruskal-Wallis test. Dunn’s Multiple Comparison post-hoc test with Bonferroni correction was used to determine the MIC of each treatment against all three bacteria. We defined the MIC as the lowest treatment concentration that was significantly different from the solvent control. Propolis dilutions one through five were ultimately excluded from analysis, as these wells were oversaturated, causing noise in OD readings. All statistics were performed in R v4.0.3 (R Core Team, 2019).

## Results

Growth of *M. plutonius* was inhibited by OTC (p < 0.001) and all three hive product extracts (Fig. 2A). The MIC of honey comb extract (p = 0.007) and brood comb extract (p < 0.001) was 16.00 mg/mL and 45.33 mg/mL, respectively. The MIC of propolis extract against *M. plutonius* was the second-lowest concentration tested, 1.14 mg/mL (p < 0.001). Growth of *S. saprophyticus* was inhibited only by OTC (p < 0.001) and propolis extract (Fig. 2B; p = 0.001). The MIC of propolis extract against *S. saprophyticus* was 2.28 mg/mL. Growth of *E. coli* was not significantly inhibited by any extract, but was inhibited by OTC (Fig. 2C; p = 0.001). Statistical analyses also revealed a significant effect by plate for each bacterium (p < 0.001), as well as a significant effect of extract source location for *S. saprophyticus* (p < 0.001) and *M. plutonius* (p = 0.032).

## Discussion

In this study, we set out to test the antibacterial effects of ethanol extracts of three hive products— propolis, brood comb, and honey comb—against *M. plutonius*, the causative agent of EFB in honey bees. Overall, *M. plutonius* growth was most susceptible to inhibition by extracts of all three hive products. To our knowledge, this is the first study to test the antibacterial properties of propolis and wax combs against *M. plutonius*. The highest concentrations of both honey comb (16.00 mg/mL) and brood comb (45.33 mg/mL) extracts successfully inhibited the growth of *M. plutonius*, but MICs for these extracts were an order of magnitude higher than that of propolis extract (1.14 mg/mL; Fig 2A). We demonstrated strong antibacterial properties of propolis against *M. plutonius*, at roughly the same concentration as our positive control OTC (1.0 mg/mL). The gram-positive model bacterium, *S. saprophyticus*, was more susceptible than *E. coli* to inhibition by propolis extract (MIC = 2.28 mg/mL; Fig. 2B), which is consistent with previous work showing that propolis extracts have more inhibitory activity against gram-positive bacteria (Rahman et al., 2010; Ristivojević et al., 2016; Sforcin et al., 2000; Tosi et al., 1996; Velikova et al., 2000). This tolerance is likely due to the structure of gram-negative bacterial cells, which are characterized by a thin layer of peptidoglycans and an extra plasma membrane (Harrop et al., 1989); however, the composition of these outer membranes—and thus the tolerance to propolis activity—may be species-dependent (Mirzoeva et al., 1997). In line with this previous work, propolis extracts here did not inhibit *E. coli* growth.

We observed significant differences between experimental plates, as well as between materials collected from different apiary locations. These differences are likely due to a small number of replicates. Though chemical composition of propolis is dependent upon the hive location and surrounding flora, this was unlikely to have affected our experiment, as our hives were located within 200 miles of one another, with similar surrounding flora. Additionally, propolis sourced from North America, Europe and Asia is characterized by similar antibiotic compounds, including flavonoids, aromatic acids, and esters (Sforcin, 2016).

The antibiotic properties of propolis are well-established (Marcucci, 1995; Przybyłek & Karpiński, 2019), but this is not the case for properties of honey comb and brood comb. Our results suggest that antibacterial compounds are much more abundant in propolis but are also present in wax combs and can serve to inhibit the growth of *M. plutonius*. Understanding the origins of this activity could inform beekeeping practices, such as brood comb replacement.

Beeswax itself is made up of hydrocarbon chains, free fatty acids, free fatty alcohols and wax esters, but also contains plant-derived materials (e.g., nectar, pollen, resin) that are incorporated as honey bees build and fill combs (Fratini et al., 2016). Due to its lipophilic composition, beeswax captures many compounds, including pheromones and acaracides (Svečnjak et al., 2019). Therefore, we assume that antimicrobial effects of wax comb extracts are likely due to other compounds incorporated during and after comb construction. To disentangle the effects of beeswax itself from those of other compounds taken up by or incorporated into the wax, future research should focus on collecting wax scales directly from bees (as in Svečnjak et al., 2019).

Brood comb frames within a healthy colony will typically hold cells of pollen and honey along the outer edges (Schneider, 2015), but using a queen excluder (Fig. 1; a common practice for hobbyists and large-scale beekeepers) encourages honey storage to occur outside of the brood boxes. Like propolis, honey is another bee-derived substance that has long been used in human medicine (Bogdanov et al., 2008). The antimicrobial properties of honey derive from its low pH, flavonoids present in pollen and nectar, as well as enzymes originating from both plants and honey bee hypopharyngeal gland secretions (Aurongzeb & Azim, 2011). Though we collected honey comb free of honey to make our ethanol extracts, there were likely traces of honey present, and previous honey storage may have incorporated compounds, such as flavonoids, into the wax cells.

The propolis envelope that surrounds the entire nest cavity of a feral colony has been hypothesized to have several homeostatic functions, including antimicrobial protection, water-repellency and even communication (Seeley & Morse, 1976). This extensive envelope is not present in managed colonies, but propolis is incorporated into the rims of wax cells, possibly to serve a structural function (Strehle et al., 2003) or confer an antimicrobial benefit (Evans & Spivak, 2010), but these hypotheses remain relatively unexplored. The composite structure of brood combs—containing silk, frass, propolis and exogenous plant material (Hepburn & Kurstjens, 1988)—results in a darker, thicker comb than those that are used strictly for honey storage, but studies on the quantity of propolis in brood or honey combs, and whether those quantities differ between comb type, do not exist. Though there are potential benefits from propolis build-up in well-used comb, there is still concern over build-up of pesticide residues and pathogen persistence in old brood combs, which currently informs brood comb replacement practices.

### Propolis as a potential preventative or treatment for EFB

With well-established antibacterial properties, and our demonstrated activity against *M. plutonius*, propolis and propolis extracts should be further developed for control of EFB in honey bee colonies. There is now a growing body of literature supporting the benefits of propolis in honey bee immunity, and bee stocks artificially selected for higher propolis collection demonstrate increased survivorship and longevity (Nicodemo et al., 2014). Aside from breeding bees for increased propolis collection, two of the most practical ways to supplement colonies with propolis include applying propolis extract to colony walls or stimulating propolis collection through hive design (i.e., propolis traps with many cervices; Borba et al., 2015; Drescher et al., 2017). Colonies supplemented with propolis exhibit lower viral infection rates and immune gene expression (Borba et al., 2015; Drescher et al., 2017; Simone et al., 2009). As a bee-derived substance that is (1) already present in the hive, (2) continually changing in its chemical constituents (based on available flora), and (3) safe for humans when present in harvested honey (unlike OTC; Gilliam and Argauer, 1981; Matsuka and Nakamura, 1990; Sporns et al., 1986; Wilson, 1974), propolis shows promise as a sustainable preventative or curative against EFB outbreaks. Field experiments using propolis traps or extracts, as well as *in-vitro* experiments (as in Grossar et al., 2020) in which EFB-infected larvae are treated with topical or ingestible propolis extracts, are the next step to evaluating the effectiveness of propolis to manage EFB.

## Acknowledgements

Tyler Eaton sequenced and confirmed our *M. plutonius* isolate. Robert Filbrun assisted with colony placement and management at Muck Crops Agricultural Research Station. Thaddeus Ezeji, Christopher Okonkwo, and Christopher Madden contributed materials, time, and guidance in anaerobic microbiology. Larry Phelan contributed time and guidance in data analysis. Sreelakshmi Suresh assisted in data processing.

Support was provided by state and federal funds appropriated to The Ohio State University, Ohio Agricultural Research and Development Center (OHO01277 and OHO01355-MRF), as well as The Ohio State University’s SEEDS Competitive Grants Program for the College of Food, Agriculture and Environmental Science (2019-096).

---

**Table S1.**
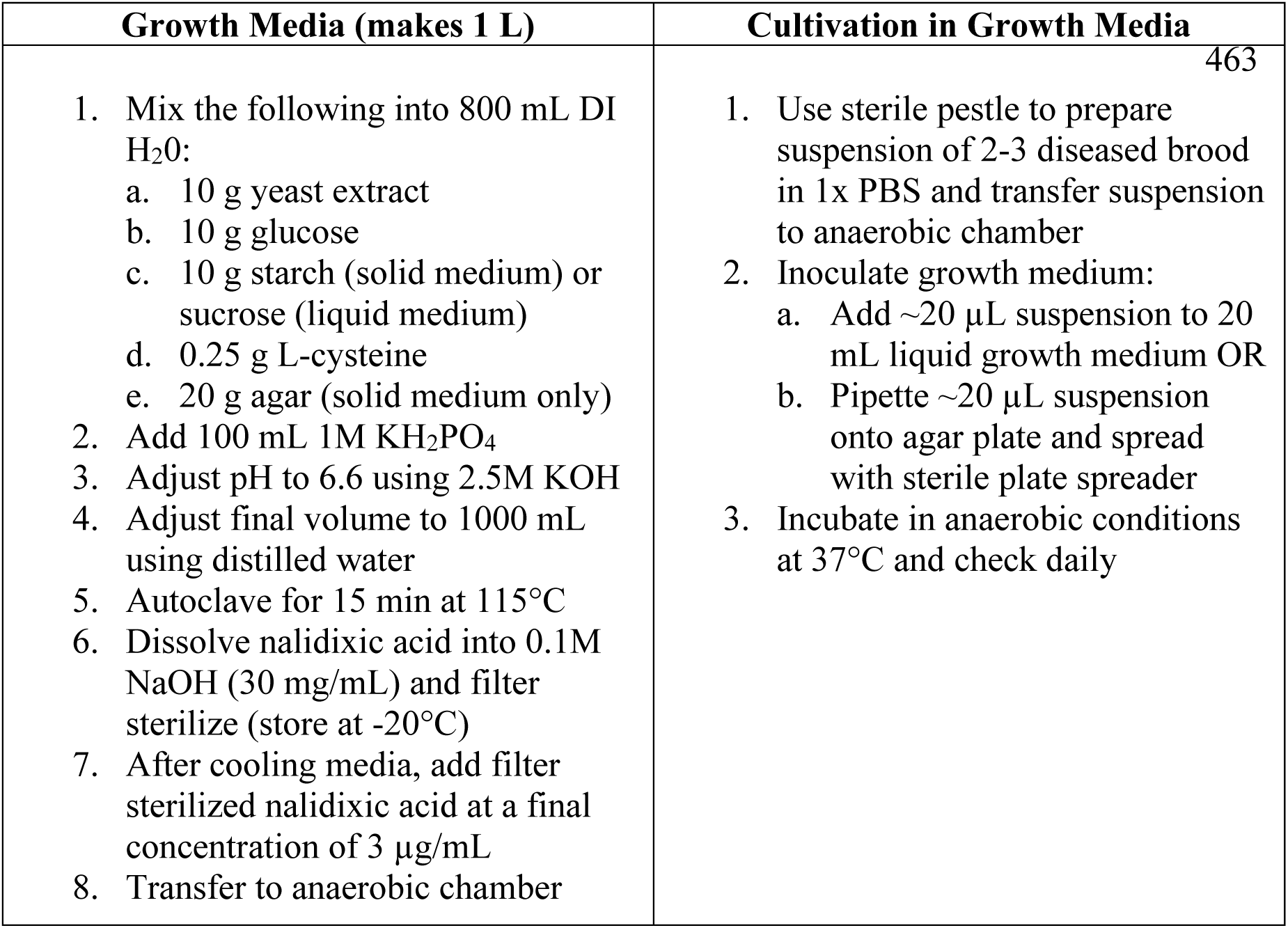
Protocol for *Melissococcus plutonius* cultivation. Growth media adapted from Bailey (1957) and Forsgren et al (2013). All steps were performed using sterile equipment and in a sterile environment, including a biosafety cabinet and an anaerobic chamber.

| Growth Media (makes 1 L) | Cultivation in Growth Media |
| --- | --- |
| Mix the following into 800 mL DI H<sub>2</sub>O:<br>10 g yeast extract<br>10 g glucose<br>10 g starch (solid medium) or sucrose (liquid medium)<br>0.25 g L-cysteine<br>20 g agar (solid medium only)<br>Add 100 mL 1M KH<sub>2</sub>PO<sub>4</sub><br>Adjust pH to 6.6 using 2.5M KOH<br>Adjust final volume to 1000 mL using distilled water<br>Autoclave for 15 min at 115°C<br>Dissolve nalidixic acid into 0.1M NaOH (30 mg/mL) and filter sterilize (store at -20°C)<br>After cooling media, add filter sterilized nalidixic acid at a final concentration of 3 µg/mL<br>Transfer to anaerobic chamber | Use sterile pestle to prepare suspension of 2-3 diseased brood in 1x PBS and transfer suspension to anaerobic chamber<br>Inoculate growth medium:<br>Add ~20 µL suspension to 20 mL liquid growth medium OR<br>Pipette ~20 µL suspension onto agar plate and spread with sterile plate spreader<br>Incubate in anaerobic conditions at 37°C and check daily |

